# Diminished gap junction coupling under diabetogenic conditions does not drive loss of functional β-cell sub-populations

**DOI:** 10.1101/2024.11.20.624571

**Authors:** Claire H Levitt, Dominic Isaacs, Maria Hansen, Vira Kravets, Jennifer K Briggs, Richard KP Benninger

## Abstract

Within the islets of Langerhans, gap junction coupling is important for synchronizing oscillatory free-calcium activity ([Ca^2+^]) and regulating pulsatile insulin release. In islets from multiple models of diabetes, gap junction coupling is disrupted, and [Ca^2+^] synchronization and pulsatile insulin is lost. Functional sub-populations have been identified within the islet that are linked to driving synchronized [Ca^2+^] and insulin release. These sub-populations can be disrupted under conditions associated with diabetes such as glucolipotoxicity and inflammatory environments, and their loss may drive islet dysfunction. Here we investigated how loss of gap junction coupling influences functional subpopulations under diabetogenic environments. We treated islets with a cocktail of pro-inflammatory cytokines, and protected gap junction coupling via co-treatment with a Cx36 peptide S293 that was previously shown to specifically prevent a decline in gap junction permeability and synchronized [Ca^2+^] dynamics. We performed calcium imaging and ChR2 stimulation, analyzed islet [Ca^2+^] dynamics and the presence of functional sub-populations including hubs and first-responders. 1h or 24h cytokine-treatment disrupted gap junction coupling, which was fully prevented by S293 peptide co-treatment. Treatment with pro-inflammatory cytokines decreased the recruitment of [Ca^2+^] upon ChR2 stimulation, increased the time between first and last responding cells upon glucose stimulation, and reduced the number and consistency of hub cells. When preserving gap junction coupling by S293 during cytokine treatment, the presence and consistency of these sub-populations was only marginally improved. We therefore concluded that while gap junction coupling is important for functional sub-populations to exert their influence on islet function, restoration of gap junctions alone is not sufficient to recover functional sub-populations upon diabetogenic conditions. Thus, preventing a disruption to intrinsic β-cell properties that define functional subpopulations is likely important for preserving these sub-populations during diabetes.

## Introduction

Diabetes Mellitus (DM) affects approximately 537 million people globally [1,2], and is characterized by the dysfunction and/or death of insulin-secreting β-cells in the islets of Langerhans in the pancreas. β-cells in the islet secrete insulin in response to elevations in blood glucose or other nutrients. At elevated glucose, β-cells show increased ATP/ADP following the metabolism of glucose, which drives membrane depolarization via closure of ATP-sensitive K+ (KATP) channels and activation of voltage gated calcium (CaV) channels. Subsequent oscillations of intracellular free-calcium ([Ca^2+^]) across the islet drive the release of insulin in discrete pulses.

The dynamics of [Ca^2+^] are synchronized across β-cells in the islet, which is facilitated by connexin-36 (Cx36) gap junctions [3–5]. Gap junctions synchronize the β-cell response to glucose despite heterogeneity in β-cell metabolism and excitability [6,7]. Under low glucose conditions, gap junction-mediated hyper-polarizing currents suppress spontaneous elevations in [Ca^2+^] from excitable β-cells [8,9]. At high glucose, gap junction-mediated depolarizing currents coordinate and enhance both first-phase [Ca^2+^] and oscillatory second phase [Ca^2+^] responses [3]. In Cx36 knockout models, synchronous [Ca^2+^] responses are disrupted [3,5,9], both first phase insulin secretion and second phase insulin pulses are diminished and glucose tolerance is impaired [5,10]. Importantly, islet gap junction coupling is disrupted under conditions associated with diabetes including by glucolipotoxicity [2,11,12] and pro-inflammatory cytokines [13] as well as in mouse models of diabetes [14–16] and human [11,12,17]. Synchronous [Ca^2+^] activity is also disrupted under conditions associated with diabetes [11,13,17,18], likely as a result of disrupted Cx36 gap junction coupling.

β-cells are heterogeneous in gene expression [19–21], glucose sensitivity [6,19], insulin secretion [22], and cell-to-cell communication [23]. As a result, β-cells exhibit functional heterogeneity, and display distinct insulin secretory patterns. This functional heterogeneity is evidenced in previous studies that found ∼75% of β-cells exhibited a fixed response to glucose, whereas ∼25% fluctuated between states of responsive and unresponsiveness [24]. The characterization of β-cell heterogeneity has led to the identification of functional subpopulations which have been shown to play distinct roles in governing the spatiotemporal [Ca^2+^] and insulin secretory dynamics of the islet [25,26]. For example, following optogenetic Channelrhodopsin-2 (ChR2) stimulation, a population of β-cells can recruit and elevate [Ca^2+^] in a large number of neighboring β-cells within the islet [27]. Similarly, ‘first-responder cells’, that are first to show elevated [Ca^2+^] upon elevated glucose stimulation, can recruit and coordinate less excitable cells to respond earlier [28]. This first-responder subpopulation is distinct from previously identified highly-connected ‘hub cells’ that exhibit a high degree of co-activity with neighboring β-cells [29] and have been shown to play a critical role in driving the synchronized oscillatory dynamics. Importantly, upon both glucolipotoxic environments and pro-inflammatory cytokines, the proportion of ‘hub cells’ declines [29]. This finding suggests that a selective loss of functional sub-populations may be sufficient to drive islet dysfunction in diabetes. However, the mechanisms underlying this loss of cells are unclear.

In this study, we sought to understand how functional sub-populations of cells are disrupted under conditions associated with diabetes. Given that gap junction coupling influences overall islet coordination, we specifically asked whether the loss of gap junction coupling that is associated with diabetes is a primary cause for the loss of these sub-populations and their influence on islet function. We first tested how functional sub-populations are influenced by pro-inflammatory cytokine treatment. We then tested a role of gap junction coupling, by using a Cx36 peptide [12] that was previously shown to specifically protect gap junction permeability and retain glucose-stimulated [Ca^2+^] dynamics under conditions associated with diabetes.

## Materials and Methods

### Mice/Animal Care

Islets expressing ChR2-YFP in β-cells were obtained by crossing Pdx^PB^-Cre (JAX #014647) and ROSA26-lox-STOP-lox-ChR2-YFP mice (JAX #024109). Islets expressing GCaMP6s in β-cells were obtained by crossing MIP-Cre^ER^ mice (JAX #024709) and ROSA26-lox-STOP-lox-GCaMP6s mice (JAX #028866). Mice were genotyped via qPCR (Transnetyx). All animal experiments were approved by the University of Colorado Institutional Animal Care and Use Committee (IACUC Protocol number: 000024). Mice were housed in a temperature-controlled environment with a 12 hr light/dark cycle and continuous access to both food and water. For mice with β-cell specific GCaMP6s expression, tamoxifen (solubilized in corn oil) was administered via IP injection for 5 consecutive days (50 mg/kg of body weight) to facilitate CreER-mediated recombination. All experiments use both male and female mice.

### Islet Isolation and Culture

Islets were isolated from mice under ketamine/xylazine anesthesia and euthanized via exsanguination. Collagenase (Type V, Sigma) was delivered to the pancreases at a concentration of 2.5 mg/mL via the bile duct. After inflation, the pancreas was surgically removed and digested (5 min at 37 °C). Islets were then hand-picked and cultured in RPMI medium (Gibco) containing 10% fetal bovine serum, 100 U/mL penicillin, 100 ug/mL streptomycin, and 11 mM glucose. Islets were incubated at 37°C, 5% CO2.

### Islet Treatment

Islets were treated overnight with a cocktail of proinflammatory cytokines (1 ng/mL TNF-α, 5 ng/mL IL-1β, and 10 ng/mL IFN-γ), which is equivalent to a ‘0.1x’ cocktail as previously used [13]. To protect gap junction coupling, islets were co-treated with 30 μM peptide S293 for 1-6 hours for experiments using optogenetic stimulation, and 24 hours for experiments measuring gap junction permeability (FRAP) and calcium activity ([Ca^2+^]).

### Fluorescent Recovery After Photobleaching and Analysis

Gap junction permeability was assessed by Fluorescence Recovery After Photobleaching (FRAP) [4]. Isolated islets were immobilized onto glass bottom dishes (MatTek) using cell tissue adhesive (CellTak, Corning) and incubated at 37°C, 5% CO2 overnight. Islets were then loaded with 12.5 mM cationic dye Rhodamine-123 (Rh123) for 30 minutes at 37°C in Hank’s Balanced Salt Solution (HBSS) at 5mM glucose. Islets were washed and imaged at room temperature in HBSS using a Zeiss LSM 800 confocal microscope with a x40 1.2NA water immersion objective. Rh123 was excited at 488 nm. Half the islet area was selected using a rectangular ROI and photobleached for 25 frames at 40% laser power, achieving ∼50% bleaching on average. Images were collected every 5 seconds for 6 to 10 minutes.

Image analysis of Rh123 dye diffusion after photobleaching was conducted using MATLAB and Image J. Fluorescent recovery was measured within a region of interest encompassing the entire photobleached area and individual cells. The initial fluorescence intensity was considered the point directly after photobleaching (IP). Fluorescence recovery was the final fluorescence (IF) once the diffusion of dye reached a plateau. The recovery was calculated as a percentage, relative to the initial intensity before photobleaching (I0) as follows:

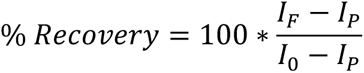

The rate of fluorescent recovery was calculated based on the linearized exponential fluorescent recovery curve of the bleached area over time (Ii) normalized to the total recovery. Calculations were conducted as follows:

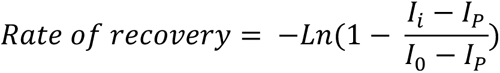

While Rhodamine 123 also loads the mitochondria, dependent on mitochondrial membrane potential, this loading does not influence the measurements of gap junction permeability [12].

### Channelrhodopsin-2 Single Cell Entrainment and Analysis

ChR2-YFP expressing islets were incubated with pro-inflammatory cytokines, with or without S293 peptide, or S293 peptide alone for 6 hours after isolation in RPMI medium (Gibco) containing 10% fetal bovine serum, 100 U/mL penicillin, 100 ug/mL streptomycin, and 11 mM glucose at 37°C, 5% CO2 prior to study.

Islets were loaded with 2μM of calcium indicator Rhodamine-2 (Rhod-2) in BMHH imaging solution (125 mM NaCl, 5.7 mM KCl, 2.5 mM CaCl2, 1.2 mM MgCl2, 10 mM HEPES, 2 mM glucose, and 0.1% BSA, pH 7.4) for 35 minutes at 37°C. Islets were then washed and imaged on a glass bottom dish (MatTek) in a 5mM glucose imaging solution using a LSM 800 confocal microscope kept at 37°C.

ChR2-YFP was excited at 488 nm. A rectangular ROI was drawn within the bounds of a single cell to avoid stimulating neighboring cells that express ChR2-YFP. The cell was activated using a photobleaching macro (intensity 5% laser power) for 8 stimulations at a frequency of 0.1 Hz. Rhod-2 was simultaneously excited at 560 nm and the corresponding Ca^2+^ response across the whole islet was measured based on fluorescent intensity of Rhod-2 that was coincident to ChR2 stimulation.

MATLAB was used to analyze ChR2 recruitment data, as previously described [27], as well as entrainment. Islets were selected by hand drawing a region of interest around the islet. Background was identified as regions that did not show any calcium response. Activity was based on the fluorescent intensity of the Ca^2+^ indicator Rhod-2 upon ChR2 stimulation and assessed using individual pixel time-courses. A pixel was considered ‘active’ if its intensity was twice that of the background. A threshold of 0.15 was used to determine which regions of the islet were activate due to ChR2 stimulation compared to the islet average. The percent of the islet area that was ‘active’ was calculated as a proportion of the entire islet area and subsequently referred to as “Area Active”. At elevated glucose, the mean Ca^2+^ time trace was averaged across the entire islet and segmented into pre and during stimulation. After detrending, a Fast Fourier Transform (FFT) was performed for both segments and the DC component subtracted. The natural oscillation frequency was identified from the pre stimulation segment as the frequency with peak power. The driving frequency was confirmed at ∼0.1Hz as the frequency with peak power during stimulation.

### GCaMP6 Imaging of Calcium Dynamics and Analysis

Isolated islets expressing β-cell specific GCaMP6 were incubated overnight with proinflammatory cytokines with or without S293 peptide, or S293 peptide alone. 30 Minutes prior to imaging, islets were fasted at 2 mM glucose in BMHH imaging solution. Islets were then transferred to a glass bottom dish and imaged on an LSM 800 confocal microscope. GCaMP6 fluorescence was excited at 488 nm with images collected at 1 frame/sec for approximately 80 minutes. 3-5 Minutes after the start of imaging, the glucose concentration was raised from 2 mM to 11 mM. For experiments measuring consistency between first responders and hub cells, glucose was subsequently lowered to 2 mM after achieving second phase oscillations for 10-15 minutes. After the islet had returned to basal activity without oscillations (∼15 minutes), the glucose concentration was increased again to 11 mM. Samples were imaged at either x20 air or x40 water immersion and held at 37°C.

Calcium dynamics were analyzed using MATLAB. First phase response time, identification of first and last responders, and characterization of highly connected hub cells was determined as previously described [28]. Briefly, to identify the response time, we determined the time at which individual β-cells reached their half maximum amplitude following stimulation from low to high glucose. This response time was rank ordered for all cells in the islet. The earliest 10% were classed as first responders and the latest 10% classed as last responders, as previously defined [29]. ΔT was calculated as the difference in time between the means of the last responders and first responders to capture the changes in spread between first and last responders between treatment groups relative to total time of the first phase. The percent consistency was determined as a proportion of top 10% of responding cells of the initial stimulation that were also first responders or nearest neighbors to first responders during repeated stimulation.

Highly correlated β-cell cells were identified using the calcium time-course at high glucose during the second phase. A highly correlated β-cell is a β-cell whose calcium activity strongly parallels many other β-cell in the islet. To quantify the degree of correlation, the degree of cross-correlation value obtained was obtained using the xcorr() function in MATLAB across all measured β-cell calcium time traces was compared among each cell. “Links” were derived using a threshold of correlation above 0.9. Hub cells have previously been defined as those possessing the “majority of connections” [29–30] from which we derived the criteria of >60% of functional “links” determined by mean degree of correlation above the defined threshold relative to all cells measured in the islet.

### Statistical analysis

#### All statistical analysis was performed in Prism (GraphPad)

A 2-sided t-test was used when comparing parameters of 2 treatment groups such as between Control versus Cytokine treated islets. Ordinary one-way ANOVA with Tukey’s *post hoc* analysis was used to compare across multiple treatment groups, where * indicates p < 0.05, ** indicates p < 0.01, *** indicates p < 0.001, **** indicates p < 0.0001. In all cases, error bars represent the standard error of the mean (S.E.M.).

## Results

### Successful protection of gap junction coupling by Cx36 S293 peptide under pro-inflammatory cytokines

To test whether functional sub-populations are disrupted by a cocktail of pro-inflammatory cytokines (0.5 ng/mL IL1-β, 1 ng/mL TNF-α, and 10 ng/mL IFN-γ), and whether sub-population loss depends on loss of gap junction coupling, we followed an approach outlined in Figure 1A. Fluorescence Recovery After Photobleaching (FRAP) was used to assess gap junction permeability and confirm a disruption to gap junction coupling by pro-inflammatory cytokines (Figure 1.B) and protection of gap junction function by the Cx36 S293 peptide, as previously described [4]. Islets treated with pro-inflammatory cytokines showed a significant reduction in gap junction permeability, as indicated by a slower recovery rate (Figure 1C, p=0.0300). During co-treatment with the S293 peptide, islets showed a 92.2%±38.6% increase in the recovery rate compared to cytokine treatment alone. No significant difference was observed between untreated control and co-treatment groups (p=0.958) or untreated control vs S293 peptide alone (p=0.918). Thus, co-treatment with S293 peptide fully protects gap junction permeability, consistent with previous studies that use the S293 peptide to protect or recover gap junction coupling in diabetogenic environments [12].

**Figure 1.**
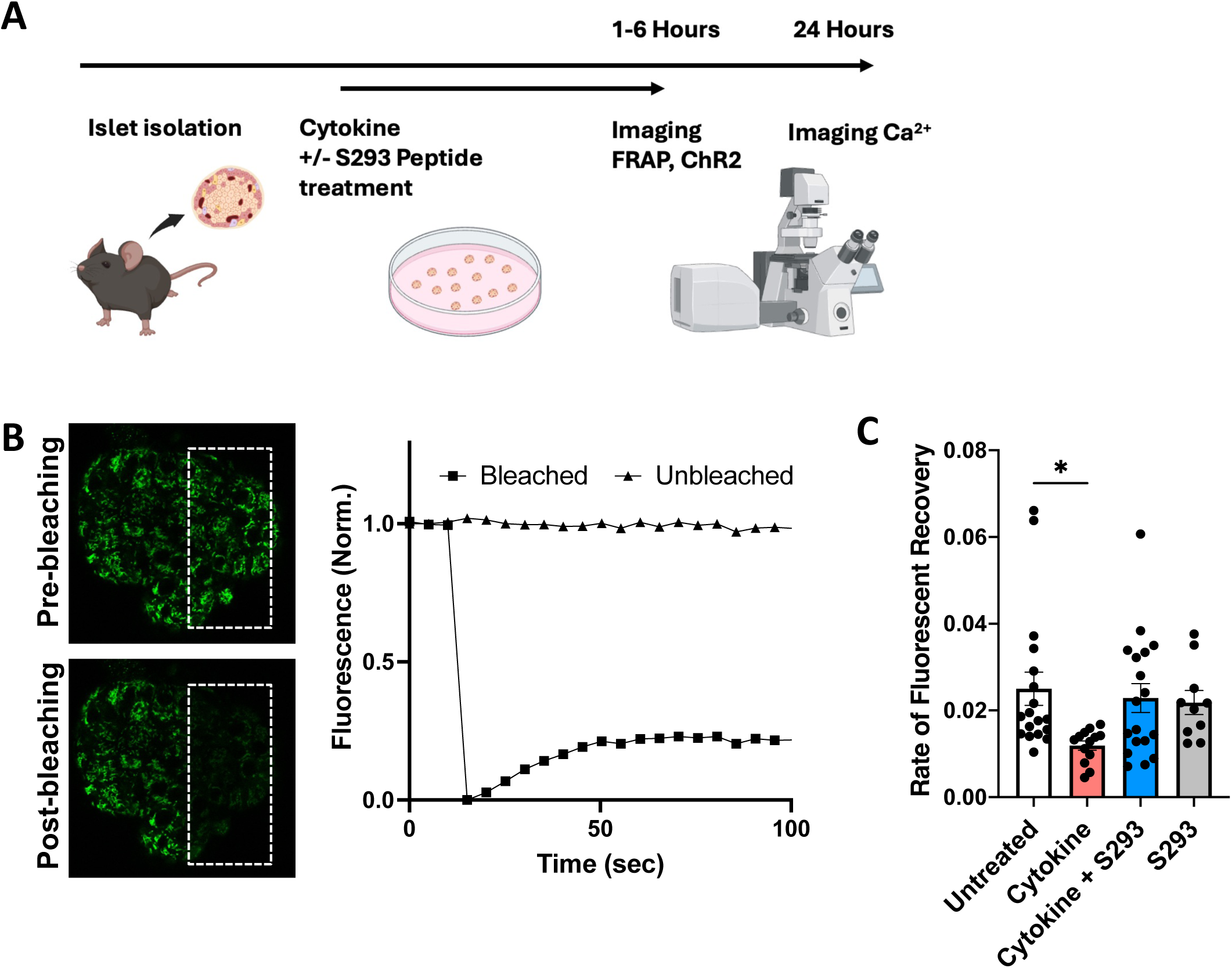
Restoration of cytokine-mediated disruptions to gap junction coupling, as measured by Fluorescence Recovery After Photobleaching (FRAP). (A) Isolated mouse islets were treated with a cocktail of pro-inflammatory cytokines (0.5 ng/mL IL1-β, 1 ng/mL TNF-α, and 10 ng/mL IFN-γ) and co-incubated with 30 μM S293 peptide for 1 hour before imaging. (B) Representative image of Rhodamine123 loaded islet pre- and post-photobleaching, along with FRAP time-course after photobleaching, showing fluorescence recovery in the bleached region. (C) Rate of recovery was calculated as a proxy for gap junction permeability in untreated, cytokine, cytokine plus S293 peptide, and S293 peptide alone treatments. Data represents Mean ± S.E.M. with each islet represented by a data point (from 6 mice), and each measurement is normalized to the mean control value per mouse. * indicates p < 0.05.

### Protecting gap junction coupling provides a partial recovery to the cytokine-mediated disruption of ChR2-mediated recruitment of [Ca^2+^] at low glucose

With the conditions established to protect gap junction coupling under a diabetogenic environment, we asked whether functional subpopulations of β-cells are also protected. Previous studies applied the optogenetic construct ChannelRhodopsin-2 (ChR2) and single cell stimulation to discover a subpopulation of β-cells with elevated metabolic activity that can recruit large numbers of neighboring cells when activated at low (5 mM) glucose levels [27]. We therefore examined how pro-inflammatory cytokines influenced these sub-populations and then tested the effects of preventing a decline in gap junction coupling on these sub-populations.

Islets from Pdx1^PB^-Cre;Rosa-LSL-ChR2-YFP mice that express ChR2 in β-cells were loaded with the Ca^2+^ dye Rhod2. ChR2 expressing cells were subjected to focal stimulation in single β-cells and the spatial Ca^2+^ response was observed in neighboring cells (Figure 2.A). We quantified the proportion of β-cells in the imaging plane that showed Ca^2+^ elevations coincident with ChR2 stimulation, which was repeated in ChR2-expressing islets treated with the cocktail of pro-inflammatory cytokines for 1 hour (Figure 2.B) and/or Cx36 S293 peptide.

**Figure 2.**
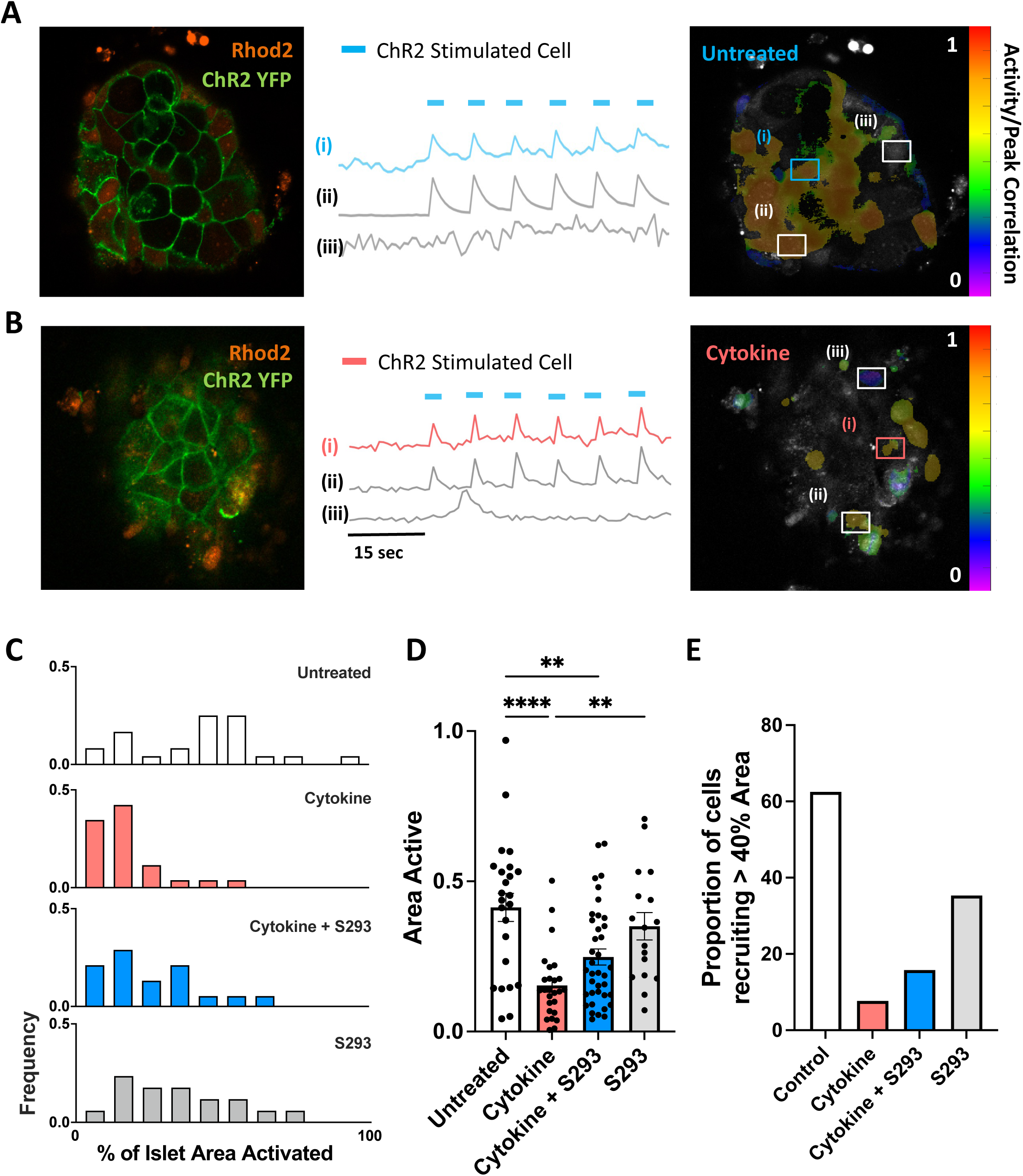
Impact of gap junction restoration on the cytokine-mediated disruption to ChR2-mediated recruitment of [Ca^2+^]. (A) A ChR2-expressing islet (left), where a single ChR2-YFP expressing cell was stimulated at 0.1 Hz (i) and calcium response was recorded across the islet which exhibited both regions of recruited (ii) and unrecruited (iii) cells in untreated conditions (middle). The result [Ca^2+^] activity was then quantified (right). (B) As in A, under cytokine-treated conditions. (C) Histogram indicating the proportion of the islet recruited by a single cell stimulation under each treatment condition. (D) Mean proportion of the islet recruited by a single cell stimulation under each treatment condition. Data represents Mean ± S.E.M. with each islet represented by a data point (from 15 mice). (E) The proportion of cells that were able to recruit an area larger than 40% of the islet under the cytokine and S293 conditions indicated. ** indicates p < 0.01, **** indicates p < 0.0001.

Under all treatments we observed heterogeneity in the recruitment of elevated [Ca^2+^] following single cell stimulation (Figure 2.C). In untreated control islets, 41.3%±4.7% of cells within the imaging plane showed elevated [Ca^2+^] following single cell stimulation, with a substantial variation (Figure 2.C,D) as previously observed [27]. Treatment with pro-inflammatory cytokines led to a significant decrease in the proportion of cells showing elevated [Ca^2+^] following single cell stimulation (15.4%±2.3%) compared to untreated controls (Figure 2.C,D, p<0.0001,). Co-treatment with the S293 peptide led to a moderate increase in elevated [Ca^2+^] following single cell stimulation (24.8%±2.6%), however this was still significantly lower than in untreated control (Figure 2.C,D, p=0.0026). Treatment with S293 peptide alone showed a similar proportion of cells with elevated [Ca^2+^] (35.0%±4.5 %) to untreated control islets (Figure 2.C,D, p=0.6680).

Untreated control islets showed broad heterogeneity in the recruitment of [Ca^2+^] (Figure 2.C), as previously observed [27]. This included a population of cells that recruited elevated [Ca^2+^] across the islet that was above the median value in untreated control islet (>40%, Figure 2.C), consistent with earlier findings [27]. Under cytokine treatment (1 hour), this ‘high-recruiting’ population of cells was almost completely absent (∼4%, Figure 2.E). Co-treatment with S293 peptide partially recovered this population of cells (∼10%, Figure 2.E), but again less than in either untreated controls or S293 peptide treatment alone. The trend in these findings were similar when we applied other thresholds for the islet area recruited upon single-cell stimulation (Supplemental 3.A).

Therefore, treatment with pro-inflammatory cytokines reduced the ChR2-mediated recruitment of cells to increase [Ca^2+^], with an abolishment of ‘high-recruiting’ cells. Protecting gap junction coupling only partially recovers ChR2-mediatred recruitment of cells to increase [Ca^2+^], with only a small number of ‘high-recruiting’ cells low glucose.

### Protecting gap junction coupling recovers cytokine-mediated disruption of ChR2-mediated recruitment of [Ca^2+^] at high glucose

Although protecting gap junction coupling did not significantly improve single-cell mediated islet recruitment at low glucose, we sought to assess whether the action of ChR2 was influenced by cytokine-mediated disruption at high glucose. Islets generally respond to changes from low to high glucose by displaying a robust first phase and sustained second phase within a 30-minute period. We therefore exposed islets to a high glucose environment for 30 minutes during which intrinsic islet [Ca^2+^] oscillations were observed, confirming that the islets had entered the second phase. We then performed single-cell stimulation of ChR2-expressing Δ-cells as we performed at low glucose (Figure 3.A-B). In untreated control islets 62.3±1.4% of the islet within the imaging plane showed elevated [Ca^2+^] that was coincident with the ChR2 stimulation following single cell stimulation at 0.1 Hz (Figure 3.C). This indicates how connected individual cells are across the islet. Treatment with pro-inflammatory cytokines led to a significant decrease in this [Ca^2+^] elevation (39.7±3.4%, p<0.01). Co-treatment with the S293 peptide led to a full recovery in elevated [Ca^2+^] (62.6±3.8%, p<0.001), indicating recovery in the overall islet connectivity.

**Figure 3.**
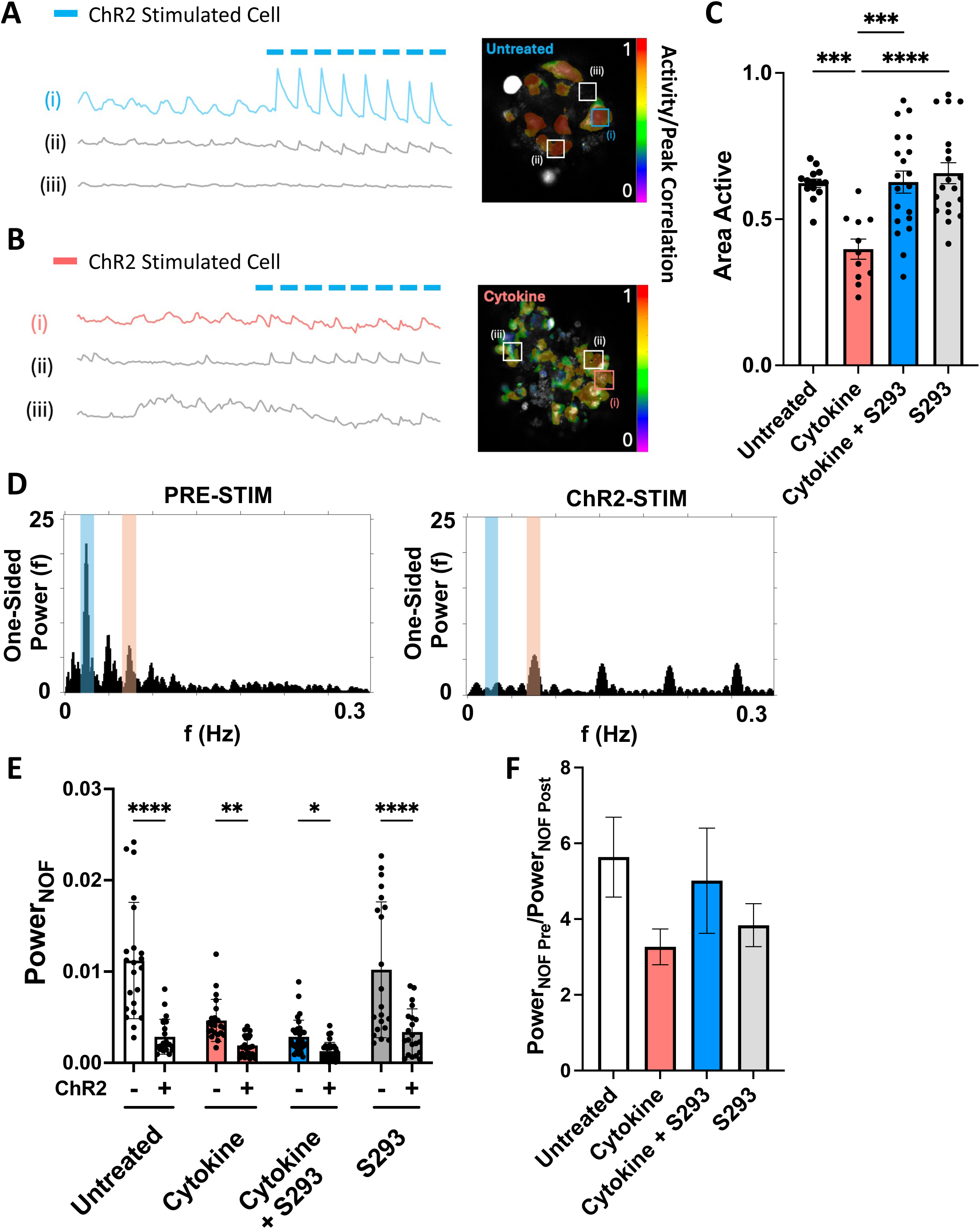
Impact of gap junction restoration *on* ChR2 single cell stimulation and islet entrainment at high glucose. (A) Representative calcium responses, where a single ChR2-YFP expressing cell was stimulated at 0.1 Hz at elevated glucose (i), and calcium response was recorded across the islet which exhibited both regions of recruited/entrained (ii) and unrecruited/not entrained (iii) cells. The result [Ca^2+^] activity was then quantified (right). (B) As in A, under cytokine-treated conditions. (C) Mean proportion of the islet recruited by a single cell at high glucose. Data presented as Mean ± S.E.M. with each islet. (D) Power spectra of the [Ca^2+^] oscillation in control islets pre-ChR2 stimulation (top) and during-stimulation (bottom). The dominant natural oscillation frequency (NOF) is highlighted in blue and the ChR2 driving frequency is highlighted in orange (E). Mean normalized power of the NOF, pre and post ChR2 stimulation under each treatment condition. (F) Ratio of the spectral power of the NOF pre/post stimulation. Data represents Mean ± S.E.M. with each experimental islet represented by a data point. All data represents n = 3 mice, 2-3 islets per condition per mouse. * indicates p < 0.05, ** indicates p < 0.01, *** indicates p < 0.001, **** indicates p < 0.0001.

We next investigated the extent to which islet cells were entrained, by analyzing the spectral power of the ‘natural oscillation frequency’ of [Ca^2+^] oscillations that are observed prior to ChR2 stimulation and analyzing the spectral power of the [Ca^2+^] oscillations at the frequency of the ChR2 stimulation (Figure 3.D, Supplemental 1.A). In untreated control islets the spectral power in the ‘natural oscillation frequency’ decreased upon single-cell ChR2 stimulation (Figure 3.E, p<0.0001). This was matched by an increase in the spectral power of the ChR2 driving frequency upon single-cell ChR2 stimulation (Supplemental 1.B, p<0.0001). Following treatment with pro-inflammatory cytokines the spectral power in the natural oscillation frequency also decreased, although this decrease was lessened compared to in control islets (Figure 3.F), indicating a less robust response to single-cell stimulation, as expected. Co-treatment with pro-inflammatory cytokines and S293 peptide increased the change in spectral power of the natural oscillation frequency compared that upon pro-inflammatory cytokines alone, and similar to that in control islets. This shows a robust recovery in the response to ChR2-mediated single-cell stimulation. Similarly, co-treatment with pro-inflammatory cytokines and S293 peptide showed a slightly increased change in spectral power at the ChR2 driving frequency, as compared to treatment with pro-inflammatory cytokines alone (Supplemental 1.C, p = 0.2274). These experiments were conducted over 10-20 islets per condition (from 3 ChR2-expressing mice), and our analysis indicates that treatment condition accounts for 19.9% of total variation indicating the observed differences in oscillation at high-glucose are attributed more to treatment rather than biological variability among samples.

Therefore, protecting gap junction coupling was able to fully recover islet-wide connectivity and entrainment at elevated glucose that resulted from ChR2 stimulation of individual cells.

### Protecting gap junction coupling does not recover the cytokine-mediated disruption to first responder cells and the 1^st^ phase [Ca^2+^] response

We next examined the impact of pro-inflammatory cytokines and the protection of gap junction coupling on first responder cells and the first phase [Ca^2+^] response. Islets from Mip-Cre^ER^;ROSA26-lox-STOP-lox-GCaMP6s mice that express GCaMP6 specifically in β-cells were used to assess changes in [Ca^2+^] following glucose elevation. We measured the [Ca^2+^] response of individual cells and ranked their response time after glucose stimulation. First responder cells were identified as the 10% of cells which showed the earliest response, and last responders as the 10% of cells that showed the latest response, with the spread in response time (Δt) being the time between first and last responders (Figure 4.A). In control untreated islets the response time was consistently short (Δt=179.5s±32.8s). Upon treatment with the pro-inflammatory cytokine cocktail for 1 hour, there was no significant impact on the time between first and last responders (Figure 4.A-B, p=0.3897). However following treatment with pro-inflammatory cytokines for 24 hours there was a significant increase in the time between first and last responders compared to untreated control islets (p=0.0043, Figure 4.A-B). Using this 24-hour treatment, co-treatment with S293 peptide did not show a decrease in Δt compared to cytokines alone (Δt=374±62s vs. 310±31s respectively), indicating protection of gap junction coupling was not sufficient to recover a narrow spread of the first phase response (Figure 4.C, p=0.72). Analysis of the response time for first responders and last responder cells alone showed that treatment with pro-inflammatory cytokines significantly increased the delay of last responding cells rather than affecting the timing of first-responding cells, consistent with a reduced first-responder action (Supplemental 4. A,B). These findings were similar when other definitions for first-responder cells were used (Supplemental 3.B). S293 peptide alone showed a similar time between first and last responders as untreated controls. Under 1 hour cytokine treatment, co-treatment with the S293 peptide had no impact (Supplemental 2.A).

**Figure 4.**
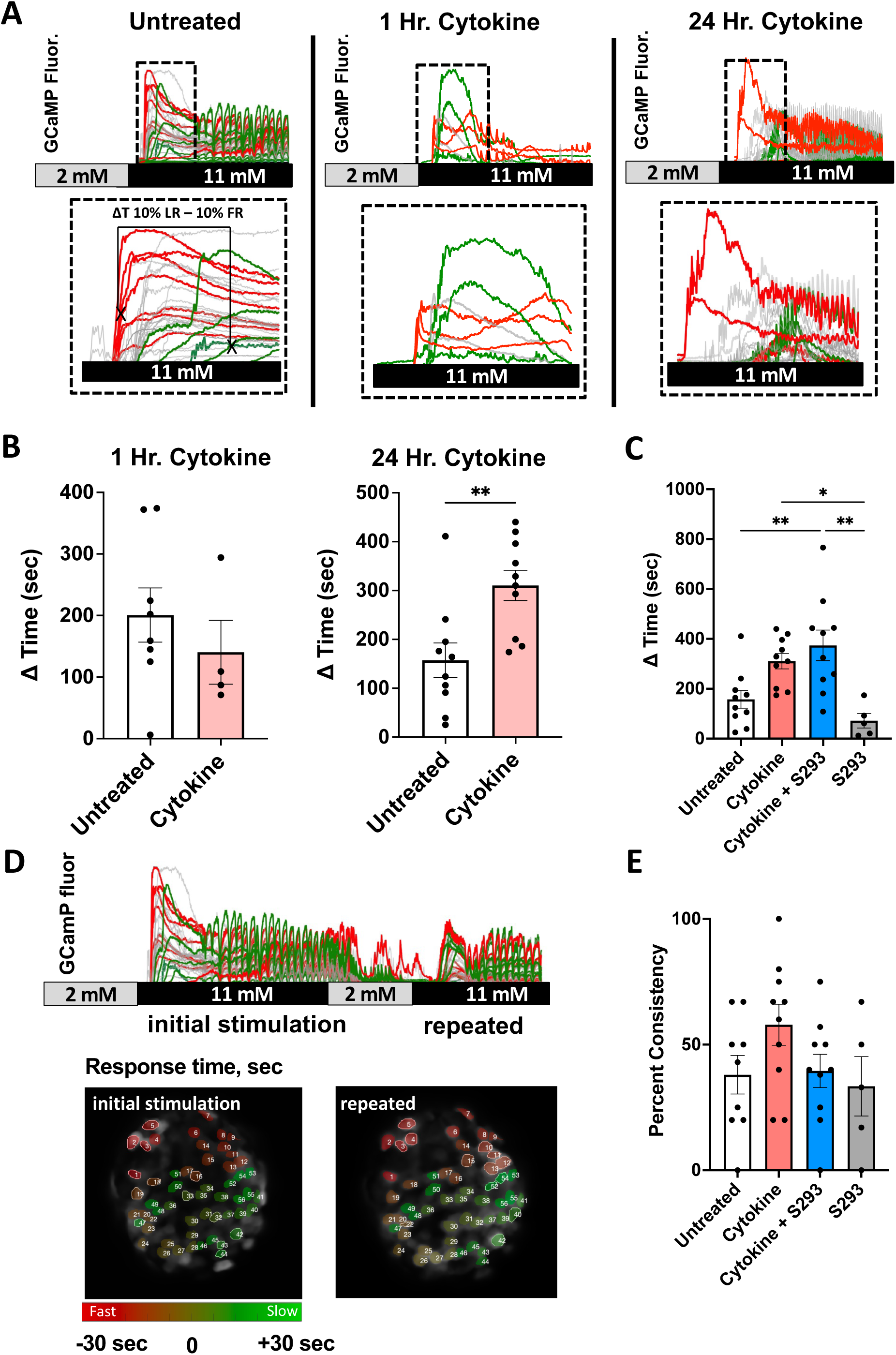
Impact of gap junction restoration on the cytokine-mediated disruption to the temporal and spatial consistency of first responders. (A) Time-course of [Ca^2+^] in first responder cells (red) and last responder cells (green) following stimulation by high (11mM) glucose, under control or cytokine-treated conditions (for 1h or 24h). (B) The mean time spread between the first and last responders (ΔT) in islets treated with cytokines for 24 hours or 1 hour, compared to untreated. Data represents Mean ± S.E.M. with each islet represented by a data point (n = 4 mice). (C) The mean time spread in islets treated for 24 hours, with cytokines and/or S293 peptide recovery of gap junction coupling. (D) Time-course of [Ca^2+^] in first responder cells (red) and last responder cells (green), subjected to repeated stimulation from low to high glucose (E) The percent consistency of first responders in untreated cells (∼40%) or those with cytokines and/or S293 peptide recovery of gap junction coupling. Data represents Mean ± S.E.M. with each islet represented by a data point (n = 9 mice). ** indicates p < 0.01, *** indicates p < 0.001.

Clusters of first responders were previously found to be consistent upon repeated glucose stimulations [28]; where ∼35% of first responder cells remained first responders and ∼75% of first responders remained within a 1 cell distance separation. We tested whether this consistency was affected by both pro-inflammatory cytokines and protection of gap junction coupling. We identified the top 10% of responding cells during an initial glucose stimulation and determined what proportion of these cells remained first responders or nearest neighbors to first responders during a repeated glucose stimulation (Figure 4.D). Upon cytokine treatment, there was a slight, but not significant, increase in the consistency of first responders upon repeated glucose stimulation compared to untreated controls (57.9% vs. 38% respectively, p = 0.2835, Figure 4.E). Upon protection of gap junction coupling via co-treatment with S293 peptide there was a slight, but not significant, decrease in the consistency of first responders relative to cytokine only treatment (39.5%, p = 0.3264, Figure 4.E), and similar to that measured in untreated controls. The consistency of first responders after 1 hour of cytokine treatment also did not significantly improve upon co-treatment with the S293 peptide (Supplemental 2.B). Therefore, treatment with pro-inflammatory cytokines disrupted the first-phase timing, which was impacted by the protection of gap junction coupling. However pro-inflammatory cytokines increased the consistency of the first phase [Ca^2+^] response, which was reversed upon protection of gap junction coupling.

### Protection of gap junction coupling provides a partial recovery to the cytokine-mediated disruption of of highly connected β-cell hubs

Finally, we examined the impact of pro-inflammatory cytokines and the protection of gap junction coupling on highly connected β-cell hubs. β-cell hubs have been demonstrated to maintain coordinated [Ca^2+^] dynamics [25,28,29], and previous work indicated a decrease in β-cell hubs upon diabetogenic conditions [13,29]. Following Ca^2+^ imaging we analyzed individual cell time-courses and formed a functional network and identified the most highly connected cells as β-cell hubs (Figure 5.A, see Methods). In control islets the number of highly synchronized functional connections and highly synchronized hubs was high and consistent with prior measurements [29,30]. Upon treatment with pro-inflammatory cytokines for 1 hour, there was no significant impact the mean degree of correlation (Figure 5.B, p=0.3708). However following treatment with pro-inflammatory cytokines for 24 hours, there was a significant decrease in the mean degree of correlation (Figure 5.B, p=0.008 and Figure 5.C, p = 0.0375). There was also a significant decrease in proportion of highly synchronized functional connections or “links” (Figure 5.D, p=0.0094) and modest decrease in highly connected β-cell hubs (Figure 5.E, p=0.3759). Co-treatment with S293 peptide did not show a significant change in neither functional connection nor β-cell hubs compared to cytokines alone (Figure 5.C-E, p=0.9832 and p=0.8101 respectively). While the proportion of hub-like cells across the entire β-cell population slightly increased upon co-treatment with the S293 peptide compared to cytokine-treated islets, this was not statistically significant. This indicates that protection of gap junction coupling was not sufficient to recover highly synchronized functional connections and highly connected β-cell hubs. S293 peptide alone showed similar functional connections and β-cell hubs as untreated controls. These findings were similar when other definitions for highly-connected cells were used (Supplemental 3.C). Under 1 hour cytokine treatment, co-treatment with S293 had no impact on mean correlation or number of links per islet (Supplemental 2.C,D).

**Figure 5.**
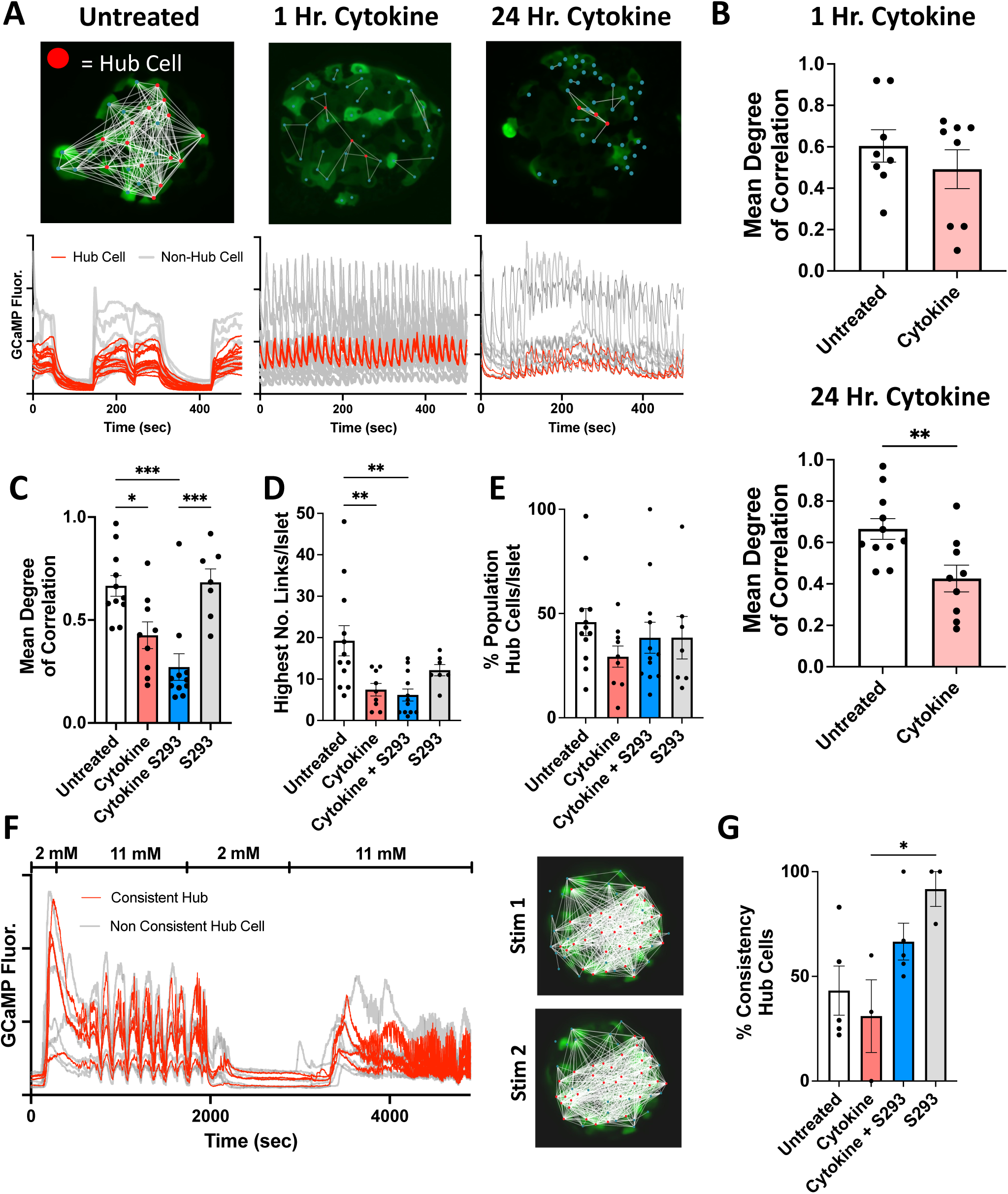
Impact of gap junction restoration on the cytokine-mediated disruption of highly connected β-cell hubs. (A) Representative connectivity map (above) and time-courses (below) of islets under control or cytokine-treated conditions (for 1h or 24h). (B) The mean correlation coefficient in islets treated with cytokines for 24 hours or 1 hour, compared to untreated. (C) Mean correlation coefficient in islets treated for 24 hours, with cytokines and/or S293 peptide recovery of gap junction coupling. (D) As in C for the average number of links between cells. (E) As in C for the proportion of hub cells among the entire islet cell population. (F) Time-course of [Ca^2+^] in β-cell hubs that are consistent or not consistent, together with representative connectivity maps. (G) The percent consistency of hubs in untreated cells (>50%) or those with cytokines and/or S293 peptide recovery of gap junction coupling. Data represents Mean ± S.E.M. with each islet represented by a data point (n = 10 mice). * indicates p < 0.05, ** indicates p < 0.01, *** indicates p < 0.001.

Approximately 43% of β-cell hubs remained as hubs following repeated glucose stimulation under control conditions (Figure 5.F). We tested whether this consistency is affected by both pro-inflammatory cytokines and protection of gap junction coupling. Upon cytokine treatment, there was a decrease in the consistency of hubs (31%) compared to untreated control islets. Protection of gap junction coupling via co-treatment with S293 peptide showed a trend toward improved consistency of hubs compared to cytokine treatment alone (66%, p=0.2109, Figure 5.G), with similar measurements to S293 peptide treatment alone, albeit lacking statistical significance. Under 1 hour cytokine treatment, co-treatment with S293 did not increase the population of hub cells as a percent, per islet (Supplement 2.E,F).

Therefore, treatment with pro-inflammatory cytokines reduced the presence of highly synchronized functional connections and β-cell hubs. Restoration of gap junction coupling, and subsequent islet-wide synchronization has little influence on hubs but maintains the consistency of those remaining hub cells.

## Discussion

Cx36 gap junctions have previously been identified as important regulators of β-cell coordination and islet function through the electrical coupling of neighboring β-cells. Pharmacological or genetic improvements in Cx36 gap junctions has been shown to recover [Ca^2+^] and insulin secretion responses in models of diabetes *in vitro*, and delay onset of diabetes *in vivo*, indicating a protective role that gap junctions play in islet function [12,31]. Functional sub-populations, which emerge as a result of β-cell heterogeneity, have been identified, and these sub-populations can drive the coordinated islet electrical response [27]. The goal of this study was to examine the influence of gap junctions on functional sub-populations of β-cells in a cytokine-mediated diabetogenic environment. Towards this, in the presence of a cocktail of pro-inflammatory cytokines, we protected Cx36 gap junction coupling using a peptide which we previously developed [12]. We then applied both optogenetics and calcium imaging to measure cell recruitment and calcium dynamics to identify and characterize three functional sub-populations that have previously been described [28] including first responders and hubs. Gap junction coupling could be successfully protected under pro-inflammatory conditions, and this was accompanied by recovery in islet-wide entrainment by ChR2 stimulation. However, optogenetic stimulation, and/or analysis of calcium dynamics revealed that functional subpopulations remained largely impaired. We conclude that while Cx36 gap junction coupling is critical for overall islet coordination and emergence of functional sup-populations, protection of gap junctions is not sufficient to recover the loss of these subpopulations upon cytokine-mediated disruption. Therefore cell-intrinsic properties are likely more influential in driving the impairment of functional sub-populations under diabetogenic conditions.

### Impairment of functional sub-populations under diabetogenic conditions

Increased levels of proinflammatory cytokines within the islet microenvironment, including IL1-β and TNF-α, is a key feature of diabetes and has been attributed to driving islet dysfunction [16,32,33]. Prior studies have found that functional sub-populations are also disrupted in diabetes, such as reduced numbers of β-cell hubs under both pro-inflammatory cytokines and glucolipotoxic conditions [29], reduced consistency of wave initiator/leader cells under high-fat diet conditions [15] and reduced presence of hubs and leaders in islets from human subjects with type2 diabetes [34]. In this study we extended these findings to show both the cells that recruit a large proportion of the neighbors to show elevated [Ca^2+^] upon stimulation at low glucose (‘high-recruiting’), and the response time of first responder cells that respond earliest to glucose were both disrupted by pro-inflammatory cytokines. This points to a general occurrence for functional sub-populations to be disrupted under diabetogenic conditions. Notably, each of these functional sub-population is associated with driving a specific aspect of islet electrical response, which were also disrupted under the pro-inflammatory cytokines. For example, β-cell hubs coordinate oscillatory dynamics across the islet, and the synchronization of these oscillations was disrupted by pro-inflammatory cytokines. First responders coordinate and enhance the initial first-phase response to glucose elevation, which was also disrupted by pro-inflammatory cytokines. Thus, disruption to these functional sub-populations may be contributing to islet dysfunction under diabetogenic conditions.

### Influence of gap junction disruption to functional subpopulations under diabetogenic conditions

Functional sub-populations, such as β-cell hubs, first-responders, and those that show high ChR2-recruitment of [Ca^2+^] at low glucose, all exert their influence to coordinate and recruit other β-cells via Cx36 gap junction coupling. Therefore, we hypothesized that pro-inflammatory cytokines disrupt these sub-populations and their action because of diminished gap junction coupling. By using the S293 peptide we could recover functional gap junction coupling equivalent to that in control untreated islets indicated by FRAP. However, we did not observe substantial recoveries in the presence and action of functional sub-populations when gap junctions were protected: we observed only partial recovery to ChR2-recruitment at low glucose, a slight recovery of β-cell hubs and no recovery to first responders. One simple explanation is non-specific effects of the S293 peptide, however prior studies did not observe non-specific effects on [Ca^2+^]. Further, we observed a recovery in islet-wide entrainment of [Ca^2+^] dynamics upon ChR2 stimulation at elevated glucose, which is expected to be more dependent on gap junction electrical coupling and thus confirms the effects of the S293 peptide. These findings therefore indicate that the disruptions to functional sub-populations under pro-inflammatory cytokines are not primarily due to disruptions to gap junction coupling.

While initially unexpected, our findings are consistent with the fact that gap junction coupling is not per se a defining feature of these sub-populations. For example, cells with high ChR2-recruitment at low glucose do not show increased gap junction coupling [27], β-cell hubs do not show increased gap junction coupling [30], and first responders show lower gap junction coupling [28]. Thus, gap junction coupling does not drive the formation of these sub-populations, rather they are defined by cell-intrinsic properties such as elevated glucose metabolism or electrical differences, as well as other environmental properties such as the potential influence of α- or δ-cells [35]. Dysfunction to these cell intrinsic properties or altered communication with α- or δ-cells is therefore likely driving their disruption. However, as discussed above, we note that some amount of gap junction coupling will still be required for these cells to exert their influence across the islet [3].

Enhancing gap junction coupling can recover islet function in diabetogenic conditions [12,16,36]. Indeed, we observed a recovery in overall islet entrainment at elevated glucose by ChR2 stimulation. This enhanced function has also been demonstrated via both genetic and pharmacological approaches in models of diabetes *in vitro* and *in vivo*. While promising, our findings indicate that targeting gap junction coupling alone will be insufficient to fully recover islet function, given that functional β-cell sub-populations will still be disrupted. For example, caloric restriction is able to improve islet dysfunction that results from a high-fat diet; yet, while gap junction coupling recovered, wave initiators/leaders did not [15]. Inclusion of the GLP1R agonist Exendin-4 with pro-inflammatory cytokines recovered gap junction coupling and coordinated [Ca^2+^] dynamics [37], while also impacting insulin secretion pathways via cAMP signaling. Thus, ensuring intrinsic factors and/or communication with α- and δ-cells are recovered will likely be required. Future work should examine which of these factors is important to both define functional sub-populations and preserve their action on islet function under diabetogenic conditions.

### Limitations and Future Work

Our findings indicate that protection of gap junctions alone is not sufficient to recover cytokine-mediated loss of functional subpopulations. However, other factors within the islet microenvironment during diabetes can also impact islet function, including gluco- and lipotoxicity, amyloid plaque formation, and other pro-inflammatory cytokines such as IL6. Thus, extending our findings to other invitro treatments and mouse models of diabetes will be needed in future work. Another important factor is that our study examined a 24-hour treatment with pro-inflammatory cytokines and/or S293 peptide. This compares to a 1-6 hour treatment time in previous studies that support the recovery of gap junctions and islet function by S293 [12]. While we observed a protection of gap junction coupling, it is possible that under more acute treatments we will observe some recovery in functional sub-populations and a further enhancement in gap junction coupling, given that Cx36 turnover has a half-life of ∼3 hours [38]. After 24 hours of cytokine treatment, observed loss of functional subpopulations may therefore result from effects of prolonged cytokine exposure in addition to disrupted gap junctions, further supporting our conclusion that recovery of gap junctions alone is not sufficient in restoring functional subpopulations under diabetogenic conditions. Nevertheless, a longer-term treatment is likely more reflective of the islet microenvironment in diabetes and should be further studied. Lastly, we acknowledge that our study primarily focuses on β-cells despite potential influences of other existing cell types including α- and δ-cells on β-cell behavior. Further studies addressing intercellular crosstalk is necessary to fully understand the global effects of cytokine treatment and the role of gap junctions.

### Summary

Overall, we demonstrate that proinflammatory cytokines disrupt the presence and action of functional β-cell sub-populations such as hubs and first responders, as well as the cell-recruitment of [Ca^2+^]. Preventing the decline in gap junction coupling was not sufficient for recovering these functional β-cell sub-populations. This correlates with gap junction coupling not being a factor that defines β-cell heterogeneity within the islet, but rather cell intrinsic properties or other aspects of intra-islet communication instead being disrupted by the pro-inflammatory environment.

---

**Supplement 1.**
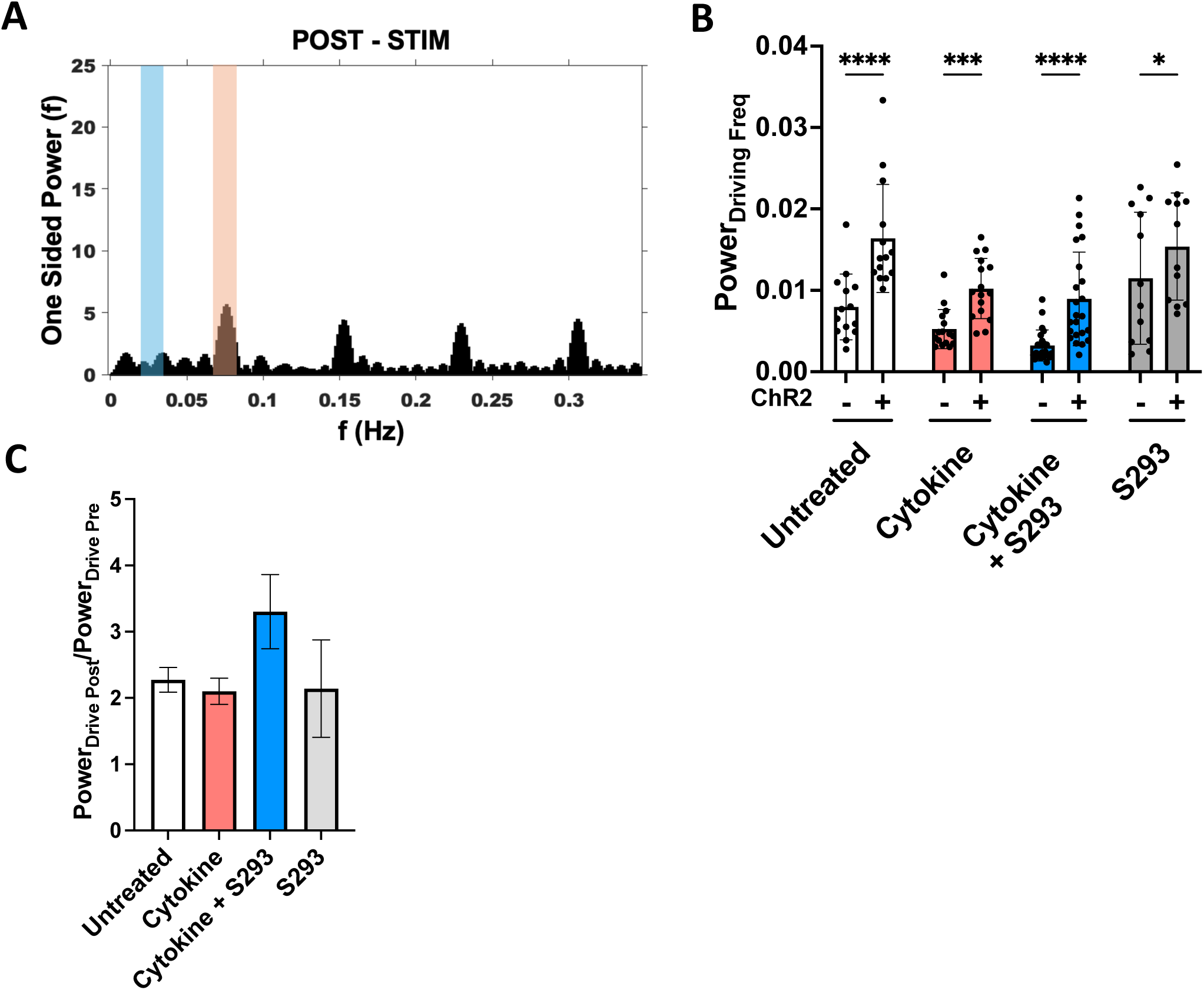
Spectral analysis of ChR2 single cell stimulation at high glucose. (A) Power spectra of the [Ca^2+^] oscillation in control islets post-ChR2 stimulation. The natural oscillation frequency (NOF) is highlighted in blue and the ChR2 driving frequency is highlighted in orange (B). Mean normalized power of the ChR2 driving frequency, pre and post ChR2 stimulation under each treatment condition. Data presented as Mean ± S.E.M. with each experimental islet represented by a data point. (C) Ratio of the spectral power of the ChR2 driving frequency post/pre stimulation averaged across all islets. Data presented as Mean ± S.E.M. with each experimental islet represented by a data point. All data represents n = 3 mice, 2-3 islets per condition per mouse. * indicates p < 0.05, ** indicates p < 0.01, *** indicates p < 0.001, **** indicates p < 0.0001.

**Supplement 2.**
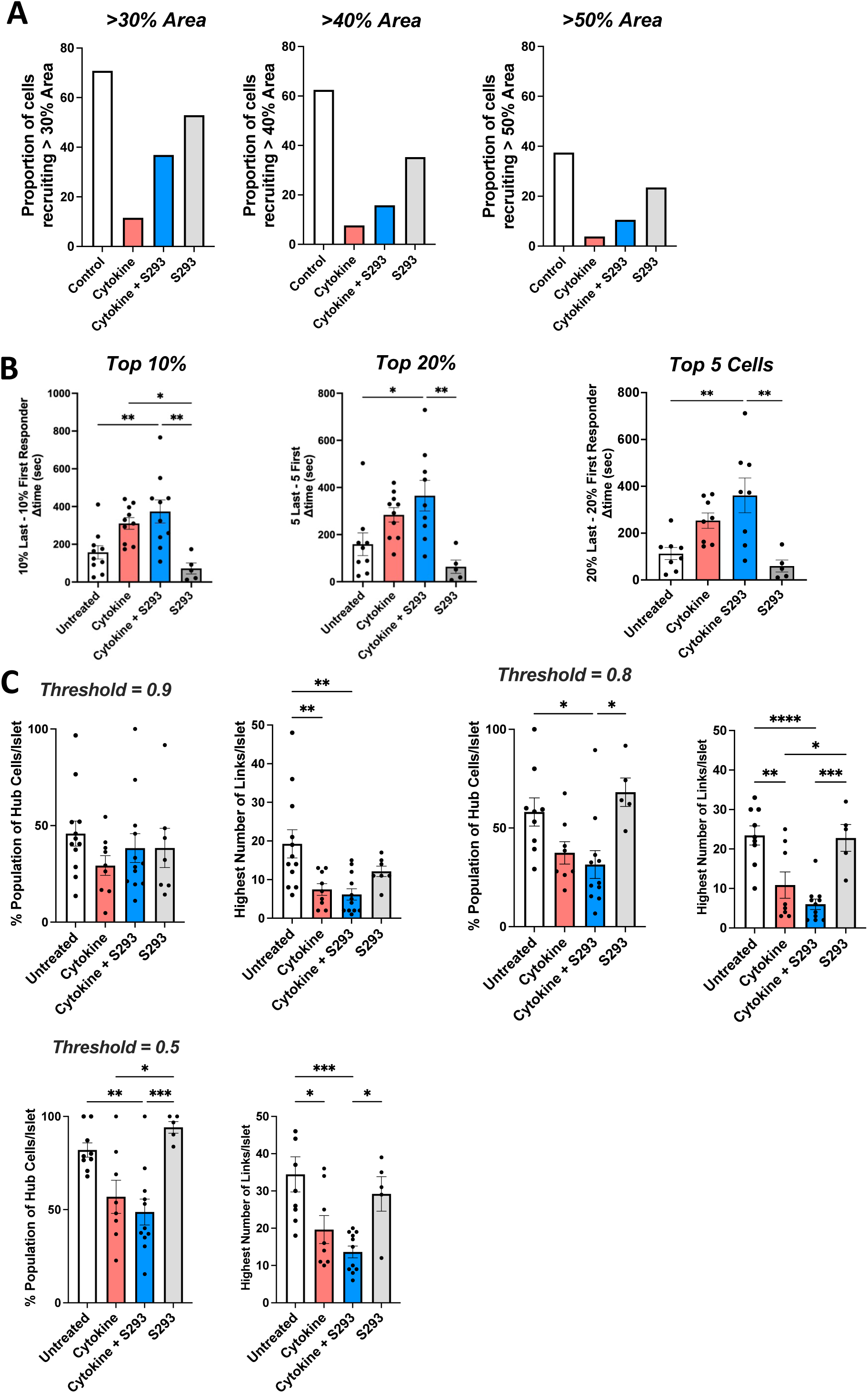
First responder and hub cells under 1h cytokine treatment with/without gap junction restoration. (A) The mean time spread in islets treated for 1 hour, with cytokines and/or S293 peptide recovery of gap junction coupling. (B) The percent consistency of first responders in untreated cells (∼40%) or those with cytokines and/or S293 peptide recovery of gap junction coupling. (C) Mean correlation coefficient in islets treated for 1 hour, with cytokines and/or S293 peptide recovery of gap junction coupling. (D) As in C for the average number of links between cells. (E) As in C for the proportion of hub cells among the entire islet cell population. (F) As in C for percent consistency of hubs. Data represents Mean ± S.E.M. with each islet represented by a data point (n = 4 mice).

**Supplement 3.**
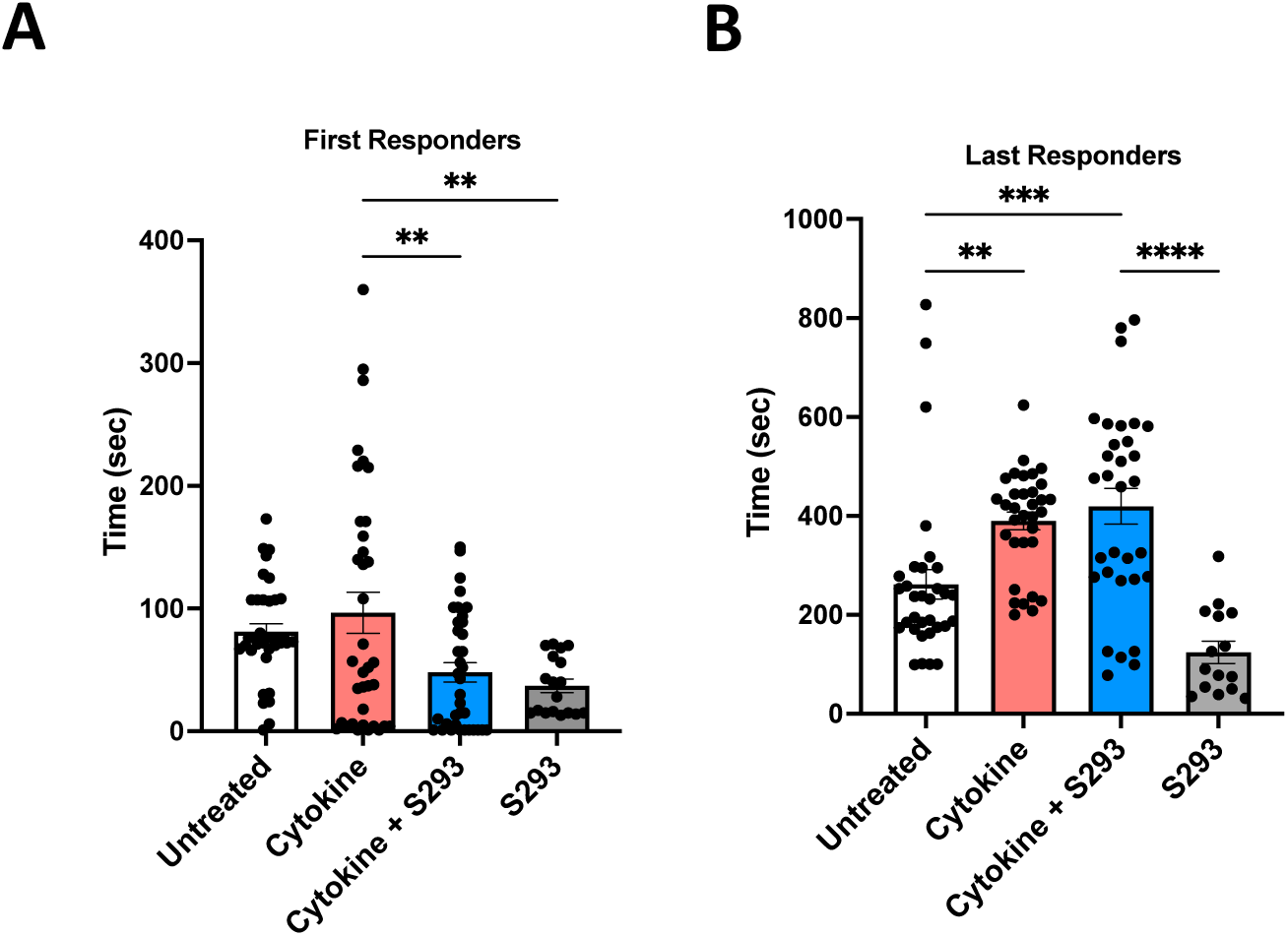
Threshold-dependent variation in defined subpopulations. (A) The proportion of cells that were able to recruit an area larger than 30%, 40% and 50% of the islet upon single cell ChR2-stimulation. Data represents mean area across all islets per condition. (B) The mean time spread between the first and last responders (ΔT) defined as the earliest and latest 10% of cells, earliest and latest 20% of cells and earliest 5 cells and latest 5 cells. (C) Population of hub cells per islet and corresponding highest number of links for correlation thresholds of 0.9, 0.8, and 0.5. Data represents Mean ± S.E.M. with each islet represented by a data point (n = 4 mice).

**Supplement 4.**
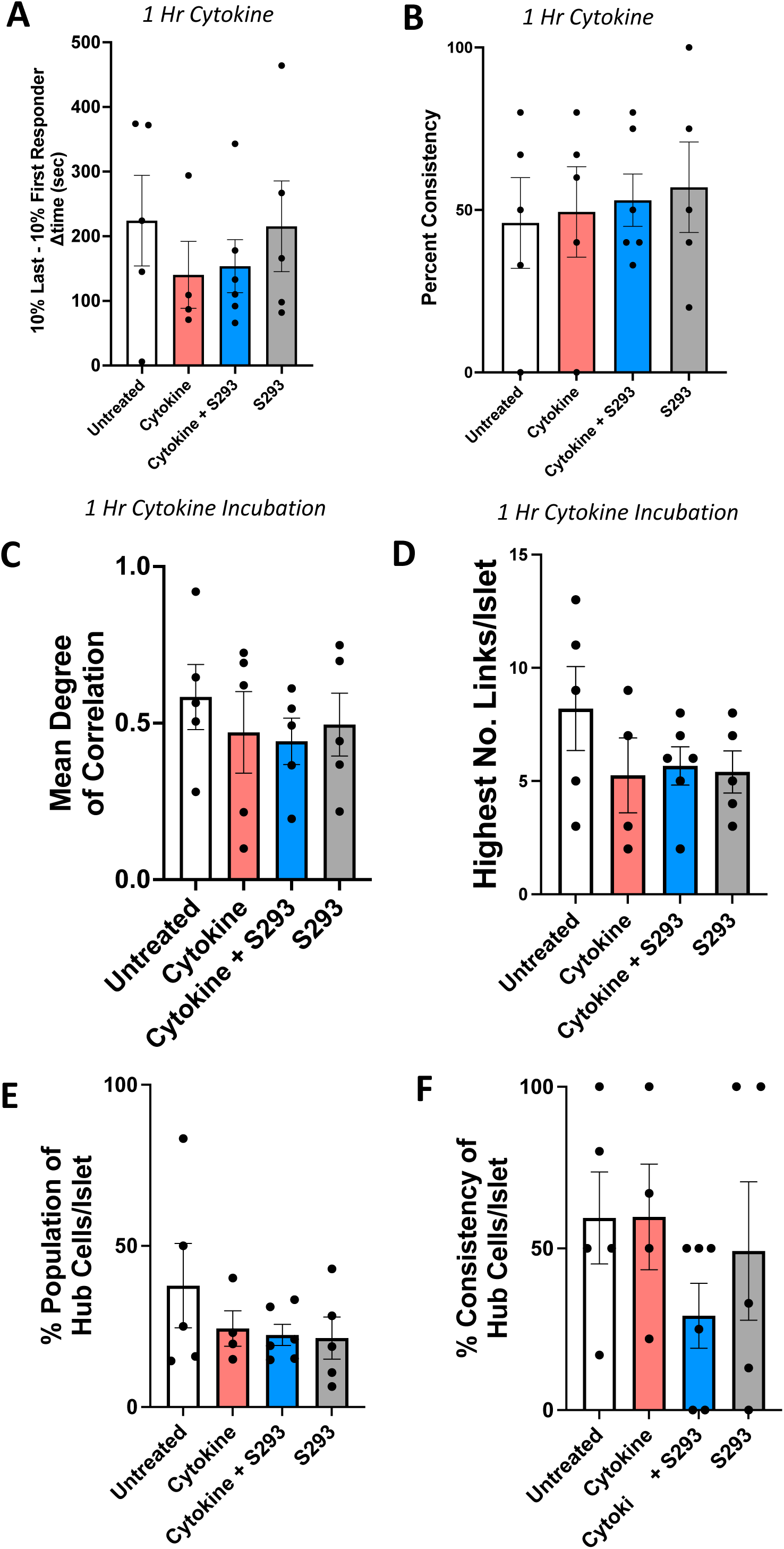
Time of first and last responder cells after 24h cytokine treatment with/without gap junction restoration. (A) The exact time of the 10% of fastest responding β-cells to reach first phase after stimulation with high glucose (B) The exact time of the 10% of slowest responding β-cells to reach first phase after stimulation with high glucose. Data represents Mean ± S.E.M. with each cell represented by a data point (n = 9 mice) ** indicates p < 0.01, *** indicates p < 0.001, **** indicates p < 0.0001.

